# SABER: A Multiparental Tomato Population Leveraging Wild Relative Diversity for High-Resolution QTL Mapping

**DOI:** 10.64898/2026.03.18.712672

**Authors:** Giovanni Gabelli, Leonardo Caproni, Fabio Palumbo, Anna Giulia Boni, Giulia Ferrari, Lucia Prazzoli, Marina Malatrasi, Sara Sestili, Matteo dell’Acqua, Massimiliano Beretta, Gianni Barcaccia

## Abstract

The narrow genetic base of cultivated tomato (*Solanum lycopersicum* L.) represents a major constraint on crop improvement, necessitating the exploitation of wild relatives to broaden allelic diversity. Here we present SABER (*Solanum lycopersicum* Allele Biodiversity Enriched Resources), a novel eight-founder Multiparent Advanced Generation Intercross (MAGIC) population that, for the first time, incorporates the Galápagos wild relative *Solanum cheesmaniae* as a founder alongside seven elite *S. lycopersicum* lines. Following a structured crossing scheme and Single Seed Descent advancement, F6 recombinant inbred lines were genotyped at 5,850 high-confidence SNP markers using Single Primer Enrichment Technology (SPET). Population structure analyses confirmed low residual heterozygosity, limited substructure among offspring, and successful introgression of *S. cheesmaniae* alleles across all twelve chromosomes. Mapping performance was validated through three Mendelian traits with known genetic determinants, all of which resolved to genomic positions consistent with the literature. QTL mapping for quantitative agronomic traits identified known loci for fruit epicarp and flesh color, and two novel QTL for days to flowering, number of leaves before flowering, and soluble solids content. Together, these results demonstrate that SABER is a powerful and reliable platform for high-resolution QTL mapping and candidate gene discovery, and establish a replicable framework for integrating wild germplasm into multiparental tomato breeding resources

## 1. Introduction

Tomato (S*olanum lycopersicum L.*) stands as one of the most economically significant vegetable crops globally, with annual production figures exceeding 182 million tons [1] Beyond its commercial dominance, the tomato is the premier model species for studying fleshy fruit development due to its well-documented physiology and the vast array of genetic and genomic tools available to researchers [2,3]. Despite this status, the genetic diversity of modern cultivars is remarkably narrow, having undergone severe bottlenecks during domestication and subsequent intensive breeding [3–5]. Historically, restricted genetic variation has hindered crop improvement, driving the need to explore wild relatives as essential reservoirs for diversifying the *S. lycopersicum* genome. While previous decades focused on utilizing wild species as sources of resilience through single-trait introgressions [1], the paradigm has recently shifted. Current research prioritizes deciphering the complex regulatory networks underlying agronomic traits and mapping the vast allelic diversity within wild populations. The recent production of pangenomes and super-pangenomes comprising wild and cultivated tomatoes exemplifies this transition toward a more holistic genomic approach [5–10].

Nevertheless, a thorough investigation into the actual agronomic impact of wild traits and alleles is still necessary. Consequently, the development of mapping populations remains an essential tool for identifying and validating valuable traits in the field. While traditional mapping approaches like biparental crosses or Genome-Wide Association Studies (GWAS) have identified major genes, they are often limited by low mapping resolution or population substructure [2]. Multiparent Advanced Generation Intercross (MAGIC) populations have emerged as powerful advanced resources to address both constraints, providing increased genetic diversity and high recombination rates that facilitate high-precision QTL mapping and candidate gene identification [3,11,12]. To date, tomato MAGIC resources have included élite lines and cherry types belonging to *S. lycopersicum* [2] or generated with interspecific crosses including the well-known *S. pimpinellifolium* and *lycopersicum* var. *cerasiforme* [3]. These studies **facilitated** the identification of valuable novel QTLs. In particular, Arrones et al. [3] demonstrated how an interspecific MAGIC population enabled the identification of key agronomic traits, such as fruit weight, locule number, anthocyanin content, leaf lobing, and the number of leaves below the first inflorescence. The extensive genetic diversity observed among wild tomato species [8,10] highlights the potential of similar studies utilizing other wild relatives.

The newly developed MAGIC population, called SABER (***S****olanum lycopersicum* **A**llele **B**iodiversity **E**nriched **R**esources), represents a significant innovation in tomato genetics by specifically targeting the untapped potential of the wild relative *Solanum cheesmaniae*. This species is highly valued for its tolerance to biotic and abiotic stresses and plant resiliency [3,13]. The SABER design introgresses *S. cheesmaniae* as a founder together with seven *S. lycopersicum* elite lines, developed for commercial, breeding and research purposes, placing these unique allelic diversity within a multi-parental framework aimed at producing innovative materials. The SABER population offers an platform for high-precision mapping of complex quantitative traits of agronomic relevance like precocity of flowering and sugar and organic acid levels in the fruit (°Brix), untapping allelic diversity that was previously inaccessible. Indeed, SABER also serves as a tool for integrating exotic diversity into modern tomato breeding pipelines, potentially contributing to cultivars better suited to challenges posed by the changing climate.

## 2. Materials and Methods

### 2.1 Plant material and founder lines

A tomato (*Solanum lycopersicum* L.) Multi-parent Advanced Generation Inter-Cross (MAGIC) population, hereafter referred to as SABER (*Solanum lycopersicum* Allele Biodiversity Enriched Resources), was developed using an eight-way design. The parental lines were selected to maximize genetic and phenotypic diversity. Seven founders belong to *S. lycopersicum* and include both elite proprietary breeding lines of Panora SpA (formerly ISI Sementi) as well as lines derived from publicly available accessions, while one founder derives from wild relative (*Solanum cheesmaniae)*.

The seven *S. lycopersicum* lines are PAR209, PAR273, PAR480, FLINE, EB5, PAR166, LA2460, the *S. cheesmaniae* line is called LA1402. A detailed description of the pedigree and the proprieties of the eight lines is in supplementary table S2.

### 2.2 Crossing scheme and population development

The SABER MAGIC population was developed following a structured eight-founder cross design aimed at maximizing recombination among parental genomes (Fig1).

**Figure 1:**
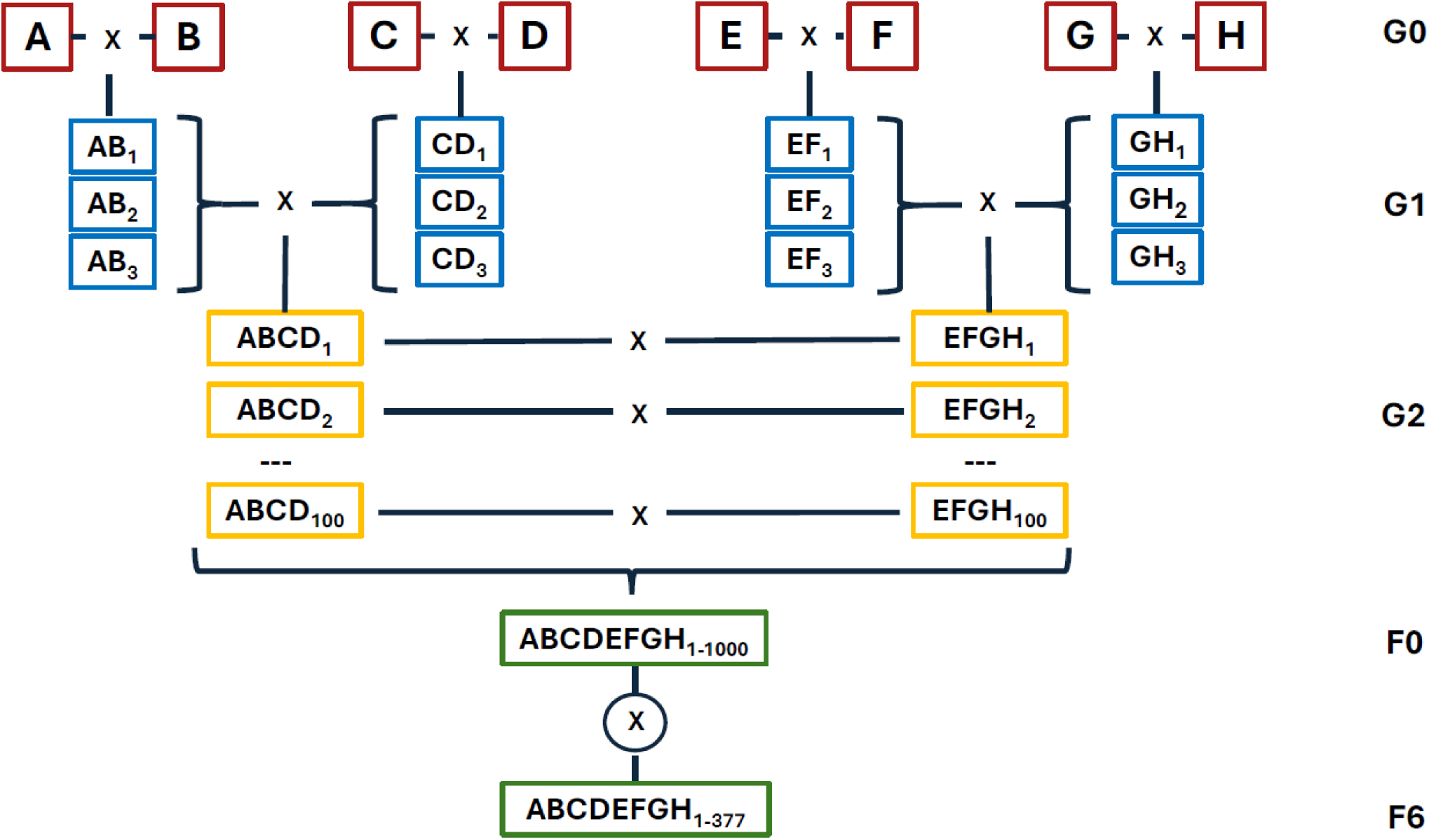
Crossing scheme of the SABER population

The 8 founder lines were initially paired to generate 4 independent two-way F₁ hybrids. All crosses involving the wild accession *Solanum cheesmaniae* LA1402, were consistently carried out using it as the pollen donor (male parent) to minimize potential fertilization barriers and improve hybrid seed set efficiency. The resulting hybrids (G1) were subsequently intercrossed as follows: the pollen of two groups of G1 was pooled and used to pollinate the individuals of the plants of the other two G1 groups, resulting in two independent four-way hybrid pools (G2). To generate the final eight-way population, 100 individual plants from each of the two four-way pools were randomly selected and grown. The same plants were crossed in a pairwise (*i.e.* one-to-one) scheme between pools, resulting in 100 independent eight-way crosses.

Seeds derived from each of the 100 independent eight-way crosses were sown, and 10 plants per cross were planted to the field, resulting in 1,000 plants representing the F0 generation of the MAGIC population. Within each progeny group (cross), 5 plants were randomly selected without phenotypic selection to maintain genetic diversity and avoid early bottleneck effects. Selected plants were advanced through successive generations using a Single Seed Descent (SSD) approach. At each generation, one seed per plant was harvested and used to establish the subsequent generation, without intentional phenotypic selection, to progressively increase homozygosity while preserving the recombination structure established during the early intercross phases. However, in cases where a progeny row displayed pronounced phenotypic variability suggestive of residual segregation, a modified SSD procedure was applied. Specifically, two individual plants from the same progeny row were independently advanced, thereby generating parallel sub-lines (“forks”). This strategy allowed the preservation of segregating genetic variation that might otherwise have been lost under strict single-line advancement, while maintaining the overall SSD framework.

### 2.3 Plant growth conditions and phenotyping material

Plants were grown under open-field conditions at Panora experimental facilities (Northern Italy, lat.44.8485154, long.10.0295428), following standard agronomic practices for determinate tomato cultivation typical of the Po Valley production system. Seeds were sown in 260-cell trays filled with sterile peat-based substrate and maintained under nursery conditions until seedlings reached the 3°–4° true leaf stage. Planting to open field was carried out approximately 40–45 days after sowing. Field planting was performed mechanically, with an in-row plant spacing of 25 cm and an inter-row distance of 140 cm. Consecutive experimental plots were separated by 100 cm to reduce border effects and facilitate management operations. Irrigation was provided with drip lines using well water. Fertigation was scheduled according to crop developmental stage and agronomic requirements. Standard crop protection practices were applied, including calendar-based fungicide treatments to prevent major fungal diseases, in accordance with integrated pest management guidelines for the region.

Young leaf tissue was sampled from each of the seedlings grown under controlled conditions (25 ± 2 °C with 80–90% relative humidity) in plastic pots filled with sterilized peat-based soil mixture. Genomic DNA was isolated using the Magnetic Bead System of the Norgen’s Plant DNA Isolation Kit (Norgen Biotek Corp, Thorold, ON, Canada) following the manufacturer’s protocol. DNA quantity and quality were verified at 260/280 and 260/230, with the NanoDrop 2000 spectrophotometer (Thermo Fisher Scientific, USA). Samples were sent to IGA Technology Services (IGATech, Udine, Italy) for high-throughput genotyping via Single Primer Enrichment Technology (SPET) [14,15]

### 2.4 Phenotyping

The phenotyping of qualitative traits (Supplementary Table S1) was carried out by observation. Obscuravenosa was assessed by observing 3 leaflets per plant, 3 plants/plot against solar light to ensure consistency in scoring. The phenotypes of the fruits (green shoulder, epicarp color, flesh color), were phenotyped by observing 3 the fruits present in the first truss of each plant, 3 plants per plot to increase the reliability of the scoring. The hypocotyl color of each accession was classified according to 3 categories: "green" (absence of anthocyanins), "purple" (presence of anthocyanins), or "segregating", observing 13 plants for in nursery containers. Days to Flowering (DTF) and leaves number before flowering (LN) were determined by observing the opening date of the first inflorescence for each plant and counting their leaves. The measurements of Soluble Solids Content (°Brix) were performed using a digital refractometer (Atago PAL-1). For each accession, the juice was squeezed from five fully mature fruits, and the average value was calculated. A total of 240 samples were phenotyped for the illustrated traits.

The qualitative phenotype data were formatted for the qtl mapping as following: green shoulder, obscuravenosa and hypocotyl color were transformed in 0/1 character to be suitable for a linear model, with segregating phenotypes assigned to NA; epicarp and flesh fruit color in 0/1/2 (yellow / orange / red) to account for an additive model, or 0/1 (yellow – orange / red). The two transformed data sets gave identical results

### 2.5 Genotyping of parent lines and SPET probes definition

Genomic libraries were prepared using the Celero™ DNA-Seq Library Preparation Kit (Tecan Genomics, Redwood City, CA, USA) according to the manufacturer’s protocol. DNA input material and the resulting libraries were quantified using a Qubit 2.0 Fluorometer (Invitrogen, Carlsbad, CA, USA). Library quality and fragment size distribution were assessed with the Agilent 2100 Bioanalyzer employing the High Sensitivity DNA assay (Agilent Technologies, Santa Clara, CA, USA). Sequencing-ready libraries were subsequently pooled and sequenced on an Illumina NovaSeq 6000 platform in paired-end mode with 150 bp read length. Raw sequencing data were processed for base calling, format conversion, and sample demultiplexing using Bcl Convert v3.9.3 (Illumina). Demultiplexed reads were aligned to the tomato reference genome (SL4.0) using BWA-MEM v0.7.17-r1188 [16], implemented in Clara Parabricks v4.4.0 with the parameters --optical-duplicate-pixel-distance 2500 and --K 1000000.

Variant discovery was performed using GATK v4.1.0.0 [17,18], following the GenomicsDBImport, ReblockGVCF, and GenotypeGVCFs workflow to identify SNPs and small insertions/deletions (indels). To minimize false positives, variants overlapping annotated repetitive regions (ITAG4.0 RepeatModeler annotation [19]) were removed using bedtools subtract v2.30.0 [20]. Subsequent quality control was performed using bcftools v1.13 and GATK [18,21]. Initial hard filtering for SNPs was applied using the criteria: QD<2.0 ∥ MQ<40.0 ∥ FS>60.0 ∥ SOR>3.0 ∥ MQRankSum<−12.5 ∥ ReadPosRankSum<−8.0, Total site depth INFO/DP<50. At the genotype level, high-confidence alternative homozygous sites were identified requiring a minimum allele depth ≥5, while maintaining sites with ≤2 missing calls and ≥5 reference or missing genotypes. For heterozygous sites, allelic balance was estimated by calculating variant allele frequencies at positions with a minimum coverage ≥10x. Finally, selected variants were converted to BED format, merged across samples using bedtools merge v2.30.0 [20] (maximum distance 20 bp), and filtered based on inter-marker distance thresholds to define the final set of 10,413 high-confidence genomic regions.

### 2.6 F6 samples genotyping

The DNA extracted from the samples was quantified using the Qubit 2.0 Fluorometer (Invitrogen, Carlsbad, CA). Libraries were prepared using the “Allegro Targeted Genotyping” protocol from NuGEN Technologies (San Carlos, CA), using 2.5 ng/μL of DNA as input and following the manufacturer’s instructions. Libraries were quantified using the Qubit 2.0 Fluorometer, and their size was checked using the High Sensitivity DNA assay from Bioanalyzer (Agilent technologies, Santa Clara, CA) or the High Sensitivity DNA assay from Caliper LabChip GX (Caliper Life Sciences, Alameda CA). Libraries were quantified through qPCR using the CFX96 Touch Real-Time PCR Detection System (Bio-Rad Laboratories, Hercules, CA) and run on the Illumina NovaSeq 6000 (Illumina, San Carlos, CA). Base calling and demultiplexing were performed with Illumina bcl2fastq v2.20. The reads of the F6 samples were processed for adapter trimming and quality filtering with fastp v0.20.1 [22], using default setting, and the resulting files checked with multiqc v1.12 [23].

The resulting reads were then aligned against the fourth release of the *Solanum lycopersicum* (SL4.0 [19]) reference genome produced by the International Tomato Genome Sequencing Project using bwa-mem2 [16] with default setting.

The SAM files of all samples were filtered for alignment quality (-q 5), sorted, and indexed with the samtools suite v1.13 [21]. A quality check was performed with Qualimap bamqc v2.3 [24]. Variant calling for all samples was performed with the GATK suite v4.1.0.0 [18] in three steps: first, variants were called for each sample using HaplotypeCaller in GVCF mode, setting the minimum base quality score to 5; second, the individual GVCF files were combined with CombineGVCFs; finally, variants were jointly genotyped for all samples using GenotypeGVCFs. Due to the size of the data, the last step (GenotypeGVCFs) was performed one chromosome at a time, and all chromosomes were merged at the end of the process with bcftools concat.

The vcf file with the variants of the founder lines was filtered with bcftools v1.13 [21] to retain only the SNPs that were homozygous in all samples, missing in none, and biallelic across the eight lines. For the F6 lines, the VCF files were filtered using custom awk commands and bcftools and vcftools v0.1.16 programs [21,25]. We retained only variants satisfying all of the following criteria: being a SNP (indels were excluded), having a mean Phred score of the conditional genotype quality across samples (GQ field in the VCF) higher than 30, having a mean of the read depth across the samples (DP fields) higher than 30, being located on an assembled chromosome (excluding SNPs on chromosome SL4.0ch00, which contains unassembled contigs), being included in the filtered variant list of the eight parent lines, having less than the 20% of the genotypes heterozygous and less of the 20% of the genotypes missing. We then identified samples with >20% missing genotypes or >10% heterozygous genotypes and removed them from the analysis. Finally, the list of variants of the parent lines absent from the list of the F6 lines were removed and vice versa, so that the two lists contain the same variants.

### 2.7 LD decay estimates and sample clustering

The filtered variant file was used in HapMap obtained by the transformation of the vcf file with plink v2.00a3 [26] to estimate the LD decay along all the genome, with a custom R script derived from Caproni et al. [27]. In brief, pairwise LD (expressed as *r^2^*) was calculated for each chromosome independently using the *LDheatmap* package [28]. LD decay as function of physical distance was estimated for each chromosome by fitting a nonlinear model based on the Hill and Weir equation. The expected *r^2^* values were computed using nonlinear least squares, assuming a population size of 254 individuals. For each chromosomethe LD half-decay distance was calculated using an overall r² value corresponding to the maximum interpolated half-decay r² (*i.e.* 0.235).

Genetic structure and diversity within the SABER were assessed using a set of SNP markers pruned for linkage disequilibrium (LD) at an r² threshold of 0.3. LD pruning was performed in plink2 using the *indep-pairwise* function with a 150-variant window and a step size of 15 variants. A version of this script is available in Macharia et al. [29]

The pruned dataset was used to estimate genetic pairwise distances among RILs using Euclidean distance with the *adegenet* package [30]. Genetic diversity, was also represented in a Neighbor-Joining (NJ) tree was constructed using the *ape* package [31]. To further investigate genetic structure, a Principal Component Analysis (PCA) was performed using the *SNPRelate* package [32].

### 2.8 Linkage map elaboration and synteny analysis

The genetic map of the SABER was obtained by anchoring the tomato MAGIC genetic positions derived from Pascual et al. [2] to the full SNP dataset on the tomato reference genome V4 (SL4.0). Missing genetic distances were subsequently interpolated using values proportional to the physical distances. In brief, the tomato MAGIC genetic map was lifted over from SL2.4 to SL4.0. The liftover was performed with CrossMap [33], which requires as input the map to be lifted over and a chain file containing the coordinate correspondences. Since a chain file linking SL2.4 and SL4.0 was not publicly available, we generated one using the Python pipeline pyOverChain [34], providing the FASTA files of both reference genome versions, downloaded from the International Tomato Genome Sequencing Project FTP repository [35]. Markers showing positional inconsistencies between physical and genetic distances were removed. Using the curated linkage map, we derived genetic positions on the SABER based on marker physical position and interpolated the missing genetic distances proportional to marker physical distances, following a script deposited by Ferguson et al. [12]. The resulting linkage map with the physical and genetic positions, as well as the chain file, are provided in the supplemental material (Data S1, S2).

### 2.9 Genotype probabilities computation and QTL mapping

The QTL mapping was entirely carried out using the R/qtl2 package v0.38 [36]. Genome reconstruction of the RILs were estimated using a Hidden Markov Model (HMM) using the function *calc_genoprob()*, using the previously described genetic map and accounting for a potential genotyping error probability of 0.002. To account for population structure and individual relatedness within the RILs, a kinship matrix was computed with the *calc_kinship* function, using the Leave One Chromosome Out (LOCO) method.

QTL mapping was performed using a linear mixed model (LMM) via the *scan1* function. The Logarithm of Odds (LOD) scores were calculated across the genome for all traits. Significance thresholds were determined through permutation testing (10,000 permutation) to control the genome-wide false discovery rate. Significance levels were set at alpha=0.05. Significant QTL peaks were identified using the *find_peaks* function. For each identified peak, the 95% Bayesian credible interval was calculated to define the QTL boundaries, and the intervals were expanded to the nearest flanking markers.

## 3. Results and discussion

### 3.1 Genotyping of the population

The design of the sequencing loci using SPET (Single Primer Enrichment Technology) [14] resulted in 10,413 primer-targeted sites for genotyping 365 F6 samples. Sequencing and alignment metrics for these samples are reported in Supplementary Material S4. After quality filtering, 5,850 SNPs and 245 samples were retained for downstream analyses. The retained samples showed low residual heterozygosity (mean: 2.47%) and low levels of missing data (mean: 2.69%). Markers exhibited a mean density of 7.53 markers per Mbp and were distributed fairly uniformly along the genome (Fig. 2), including centromeric regions, which in *S. lycopersicum* are characterized by low gene density and low recombination rates [2]. As shown in Figure 2, specific regions of increased marker density were observed, notably a small segment on chromosome 2 (approx. 8–10 Mbp) and two broader regions on chromosomes 6 and 9 (spanning 1–29 Mbp and 10–60 Mbp, respectively). To investigate the origin of this variability, we examined the distribution of SNPs with alternative genotypes among the eight founder lines (Figure S1). Across most chromosomes, the wild founder unsurprisingly exhibited the highest level of polymorphism relative to the reference genome. However, in the high-density regions on chromosomes 6 and 9, additional variability originated from other founders: PAR480 and PAR166 on chromosome 6, and LA2460 on chromosome 9. In both regions, the high number of private SNPs in these cultivars resulted in a scarcity of loci capable of distinguishing between the other lines. In other words, the set of variants has reduced discriminatory power between certain parental donors because the majority of SNPs share the same genotype across multiple lines (e.g., PAR206, PAR273, FLINE, E5, and LA2460 in the high-variant region of chromosome 6; PAR206, PAR273, PAR166, and EB5 in the central region of chromosome 9). A similar trend was partially observed in the centromeric region of chromosome 4, where two groups of founder lines showed high similarity: (PAR273, PAR166, and LA2460; PAR480, FLINE, and EB5). These trends may have a technical explanation: when one sample is genetically much more distant than the others, it can "capture" the majority of the polymorphisms during the SPET probe design process, thereby diminishing the observable differences between the remaining samples. However, the biological characteristics of cultivated tomatoes could also contribute; the genetic variability of this highly bred species is low [5], and it is common to observe conserved genomic regions across different varieties due to the selection of specific favorable traits or unknowingly shared ancestors. The high diversity spot on chromosome 2 appears to be a different phenomenon, since almost all the alternative alleles that compose the peak are private SNPs of LA1402 (*S. cheesmaniae*).

**Figure 2:**
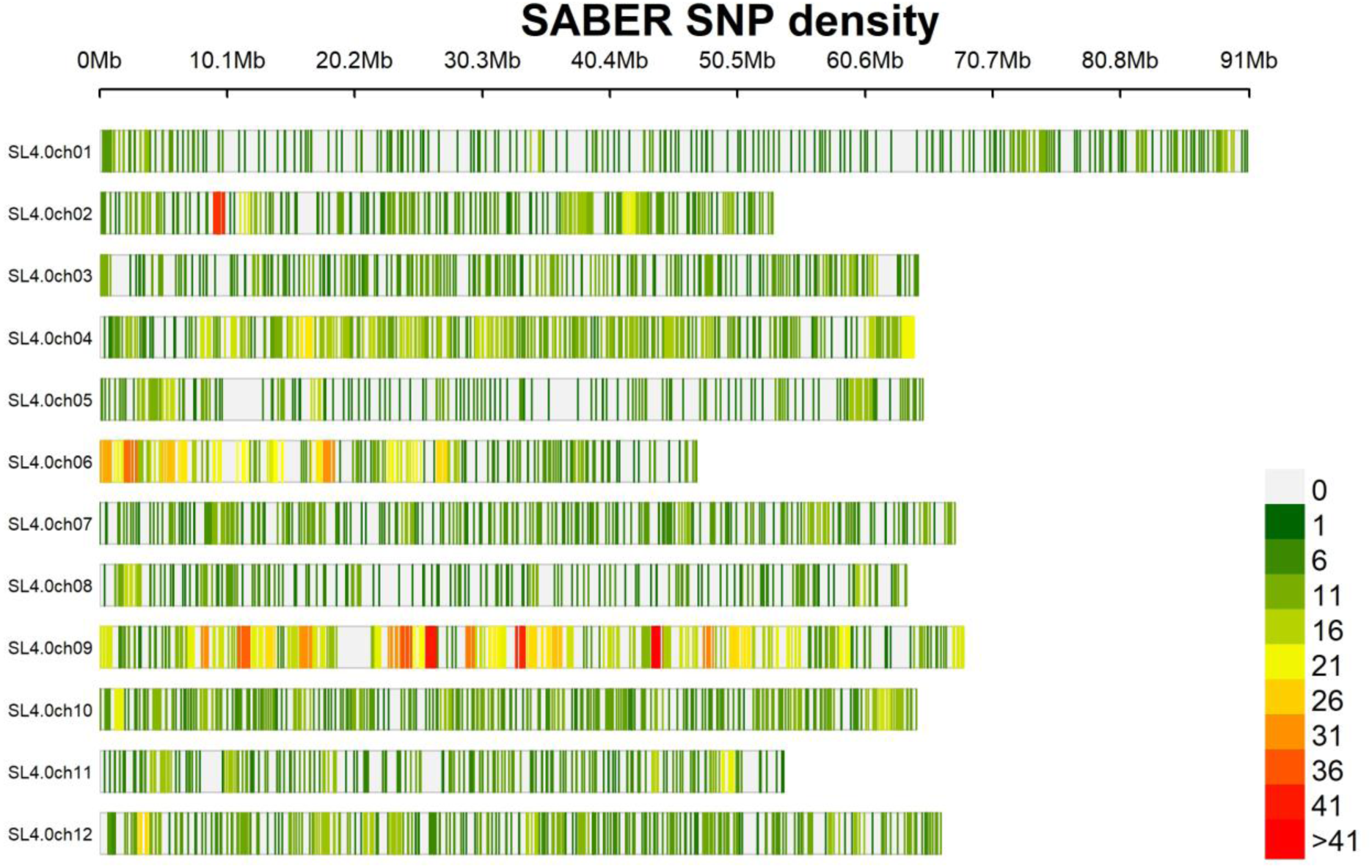
SNP density across Solanum lycopersicum genome.

### 3.2 Population structure

Principal component analysis (PCA) performed on SNP data from all lines (founders and offspring) confirmed the high genetic divergence of the wild relative compared to all other samples (Figure 3a). This pattern is particularly evident along PC1, which explains 3.03% of the variance and captures most of the diversity associated with *S. cheesmaniae*. A similar effect is observed along PC2 (2.64% of the variance), where EB5 emerges as an outlier, likely due to introgressions from *S. lycopersicum* var. *cerasiforme*. Notably, the loci differentiating LA1402 and EB5 from the other founders appear to be largely mutually exclusive. The remaining founders cluster closely together: LA2460 is genetically closest to LA1402, and FLINE to EB5, while PAR166 occupies the opposite end of the spectrum. PAR209, PAR273, and PAR480 cluster near the center of the PCA space. Offspring samples show tight clustering, indicating low population structure, with only 12 samples deviating towards LA1402 and LA2460, likely reflecting a higher genomic contribution of these founders.

**Figure 3:**
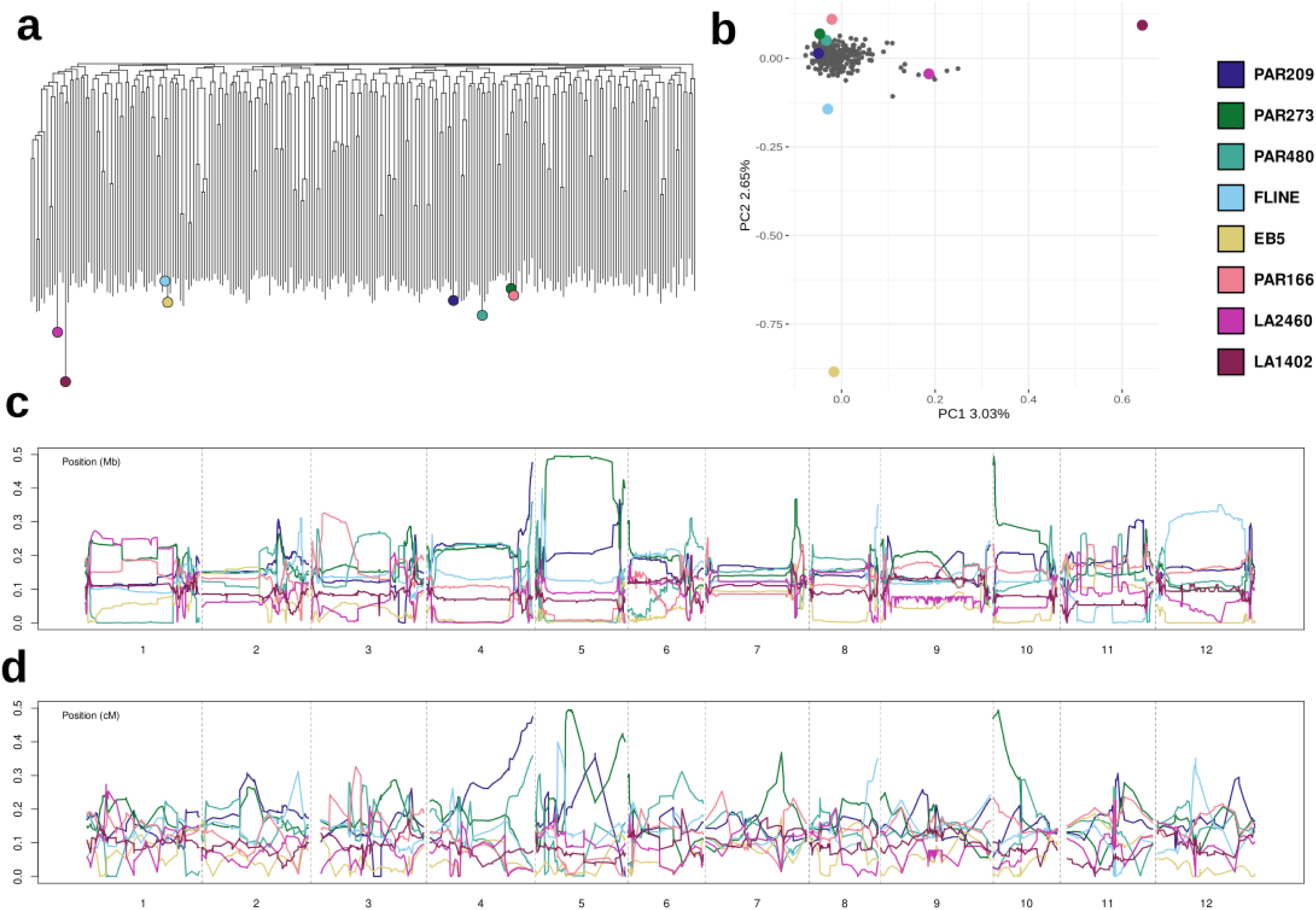
Population structure analyses of SABER population. The legend at top-right corner represents the color assigned to the eight founder lines in all panels. a) Hierarchical clustering of the SABER population. The colored dots represent the eight founder lines, b) First two principal components of a PCA of the SABER population, colored dots represent the eight founder lines c-d) Genome-wide founder haplotype blocks assignment across the 12 chromosomes according to physical (c) or genetic position (d) of the markers

Hierarchical clustering (Figure 3b) largely corroborates the PCA results, with long branches for all the samples indicating low overall population structure. Nevertheless, some pairs of samples show higher genetic similarity. This can be explained by the breeding scheme of the SABER population: the 500 RILs were selected among the 100 crosses between G3 plants (i.e. the 8-way crosses), therefore inevitably some of the remaining RILs could be descendant from full siblings.

The hierarchical clustering further confirms the wild founder LA1402 as the most genetically distinct sample, as indicated by its long branch length. PAR166 and PAR273 cluster together, consistent with their breeding history, as PAR273 served as the pollen donor in the development of PAR166. FLINE and EB5 also have a high degree of similarity in this clustering. It is unknown if these lines, both developed by public entities, share some ancestors.

The estimated average contribution of each founder to the overall population (Fig3a,b) confirms that regional imbalance is still present, but overall, the mean contribution of each parent is around the theoretical 12.5 %. A noticeable exception is represented by founder EB5, that contributes to the genome of the overall population less than expected in all chromosomes but chr3 and chr11 (Table1). The difficulties in identifying the correct donor parent in some regions due to shared genotype could help explain smaller imbalances, since they are regional phenomenon, mainly located in the large centromeric region, where few SNPs are genotyped. An example of an imbalance likely produced by technical reason is in chr5 the overperformance of PAR273 and the underperformance of PAR480 and LA2460, where all three share the genotypes in the centromere (FigS1, 3a). The low level of alleles of EB5 in the population, however, appears to have more structural causes. Since no selection has been carried out in the crossing, a random fluctuation in the first crosses could have happened. The *Solanum cheesmaniae* line, LA1402, is present in the population with levels lower than the mean, but stable across all chromosomes. This is an important result, since the interspecific hybrid could have a lower vitality. Instead, we register no regions where the contribution of the wild ancestor to the population genome is missing.

**Table 1:**
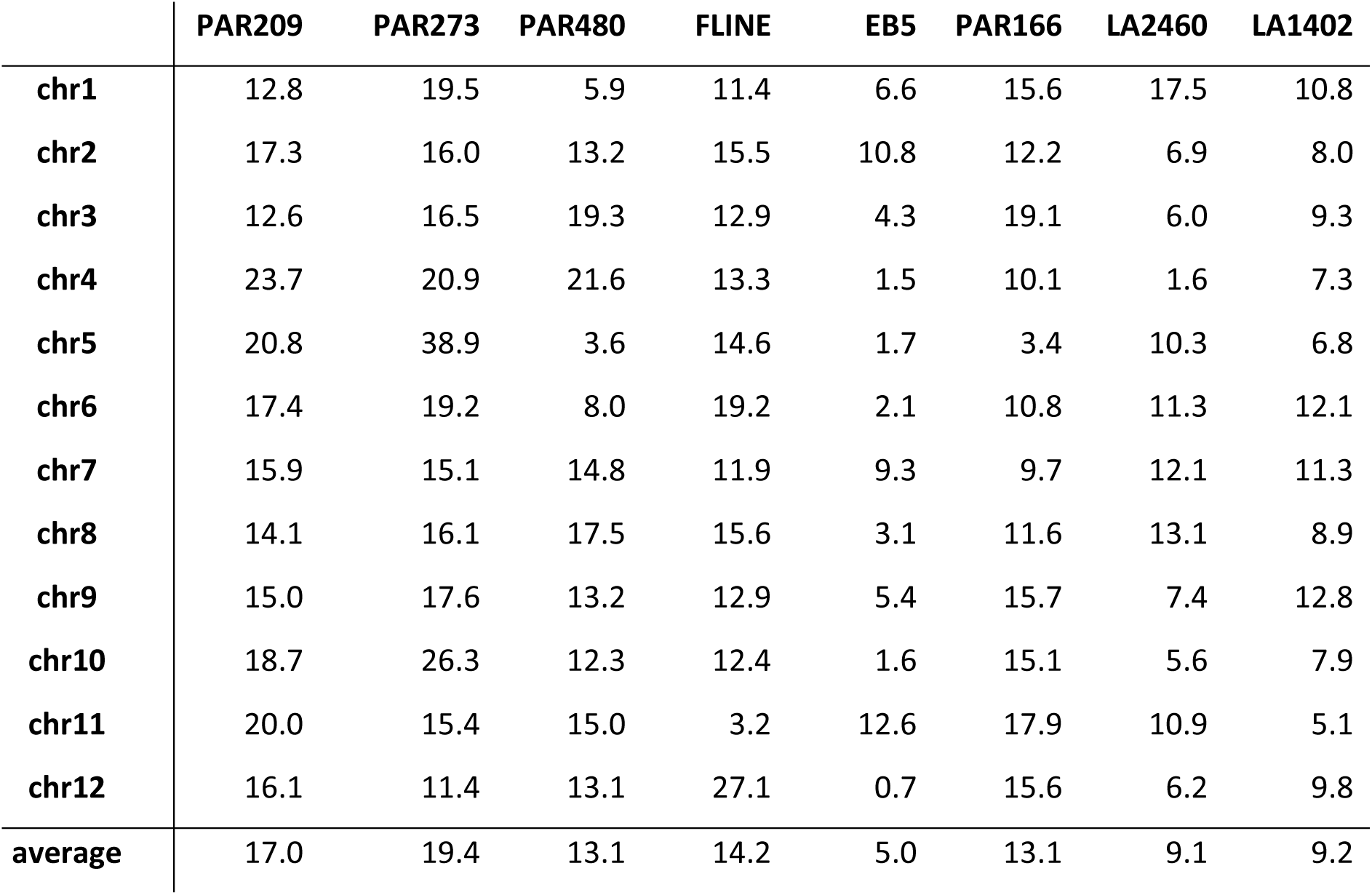
Mean contribution of each founder lines to the population genome for each chromosome. The measure is an average of the probability assigned to each marker in each sample to descend from the parent line.

|  | PAR209 | PAR273 | PAR480 | FLINE | EB5 | PAR166 | LA2460 | LA1402 |
| --- | --- | --- | --- | --- | --- | --- | --- | --- |
| chr1 | 12.8 | 19.5 | 5.9 | 11.4 | 6.6 | 15.6 | 17.5 | 10.8 |
| chr2 | 17.3 | 16.0 | 13.2 | 15.5 | 10.8 | 12.2 | 6.9 | 8.0 |
| chr3 | 12.6 | 16.5 | 19.3 | 12.9 | 4.3 | 19.1 | 6.0 | 9.3 |
| chr4 | 23.7 | 20.9 | 21.6 | 13.3 | 1.5 | 10.1 | 1.6 | 7.3 |
| chr5 | 20.8 | 38.9 | 3.6 | 14.6 | 1.7 | 3.4 | 10.3 | 6.8 |
| chr6 | 17.4 | 19.2 | 8.0 | 19.2 | 2.1 | 10.8 | 11.3 | 12.1 |
| chr7 | 15.9 | 15.1 | 14.8 | 11.9 | 9.3 | 9.7 | 12.1 | 11.3 |
| chr8 | 14.1 | 16.1 | 17.5 | 15.6 | 3.1 | 11.6 | 13.1 | 8.9 |
| chr9 | 15.0 | 17.6 | 13.2 | 12.9 | 5.4 | 15.7 | 7.4 | 12.8 |
| chr10 | 18.7 | 26.3 | 12.3 | 12.4 | 1.6 | 15.1 | 5.6 | 7.9 |
| chr11 | 20.0 | 15.4 | 15.0 | 3.2 | 12.6 | 17.9 | 10.9 | 5.1 |
| chr12 | 16.1 | 11.4 | 13.1 | 27.1 | 0.7 | 15.6 | 6.2 | 9.8 |
| average | 17.0 | 19.4 | 13.1 | 14.2 | 5.0 | 13.1 | 9.1 | 9.2 |

**Table 2:** comparison between mendelian QTLs and literature data about the genetic determinant of the mendelian traits. The distance between QTL peaks and the gene is calculated considering the nearest boundary.

| Trait | Candidate gene (SL 4.0) |  |  |  | SABER QTL |  |  |  |  |
| --- | --- | --- | --- | --- | --- | --- | --- | --- | --- |
|  | Name | Chr | Coordinates (bp) | Reference | Phenotype | Chr | coordinates (bp) | LOD | Distance (bp) |
| obv/obv+ | *Solyc05g054030* | Chr5 | 633,95,462 - 633,98,588 | Lu et al. [40] | Obscuravenosa | Chr5 | 62,575,099 | 25.1 | 823489 |
| u/ug | *Solyc10g008160* | Chr10 | 2,169,302- 2,174,039 | Powell et al. [41] | Green shoulder | Chr10 | 2,010,429 | 8.9 | 158873 |
| AH/ah | *Solyc09g065100* | Chr9 | 59,048,134- 59,055,605 | Qiu et al. [42] | Hypocotyl color | Chr9 | 59,052,650 | 29.8 | 0 |

The profiles of linkage disequilibrium (LD) levels along the chromosomes (Fig 4a) in most of the chromosomes follow the pattern of recombination frequency observed in the genetic map, with large centromeric regions with low recombination and distal region where the majority of genes are located [2,3]. Chromosomes 2 and 4 have different profiles from the others, since they are acrocentric chromosomes. Chromosomes 6, 9 and, at lesser extent, 4, exhibit high LD levels in the regions already described in this paragraph with reduced discriminating power. Indeed, this is expected since if the genotypes of two or more founder lines in a region are the same or not distinguishable, we will not detect all the recombination events that involve those lines. Considering this, and the Considering this the differences among the LD-decay levels computed along the chromosomes (Fig 4b) are not surprising, with chromosomes 4, 6, and 9 having the highest levels.

**Figure 4:**
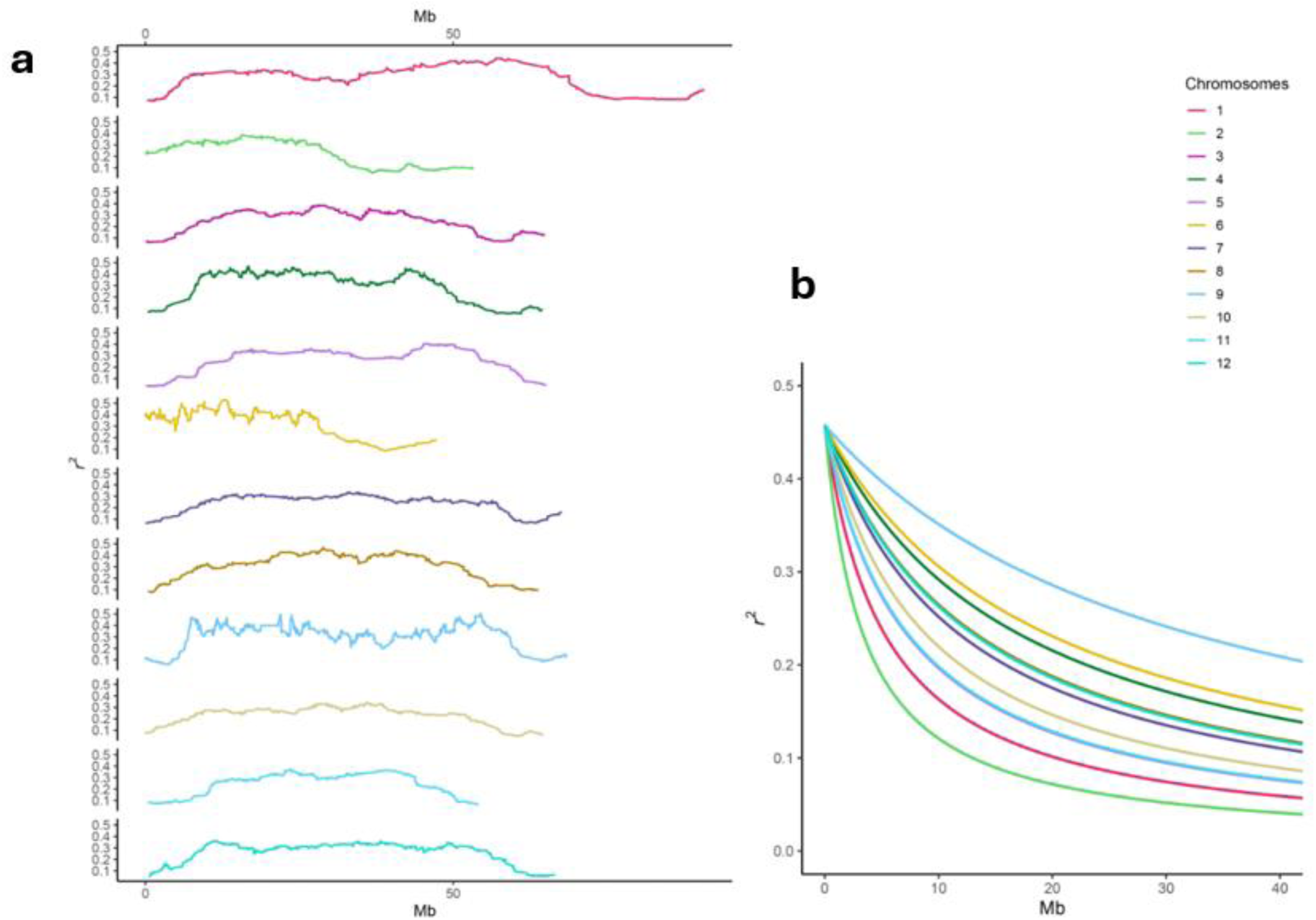
a) LD levels in the twelve chromosomes computed across sliding windows, b) LD decay in the twelve chromosomes

### 3.3. Test of mapping performances of SABER

Since the total number of SABER samples with phenotype and genotype data available is of 240, lower than the optimum recommended for similar mapping populations [37,38], three Mendelian traits with well-known genetic determinants in the literature (obscuravenosa, green shoulder, and hypocotyl color) were used to test its mapping potential.

In the majority of fresh-market tomatoes chloroplasts are not present in the subepidermal and mesophyll tissues adjacent to the leaf veins, causing the veins to appear translucent under transmitted light. In wild phenotypes and some tomato line breed for industrial processing, however, those cells contain chloroplast and veins appear of the same color for the leaf if not darker, a phenotype called obscuravenosa (*obv*) [39]. In the SABER population, a single QTL was found for this character on chromosome 5, at 62.5 Mb (LOD peak 25.1), a result that corresponds to the known genetic determinant of this character found first by Jones et al. [39], that on SL4.0 assembly is 823 Kb distant [40].

The "green shoulder" trait in tomato is a qualitative character observed during the pre-veraison stage of fruit development. In wild-type genotypes, a distinct area of intense green pigmentation develops in the pericarp tissue surrounding the peduncular insertion. The absence of this pigmentation is governed by the *uniform ripening* (*u*) mutation. While the *u* phenotype is favored in modern cultivars to ensure synchronized maturation and aesthetic uniformity for consumers, it carries significant metabolic trade-offs [41]. The QTL identified in SABER corresponds with the known location of this mutation (chr 10, 2.0 Mb, LOD peak 8.9, 159 Kb of distance from the gene).

The third Mendelian trait used to test the model is the color of the hypocotyl, that in tomato is often purple due to the accumulation of anthocyanin pigments [42]. Multiple mutants with low pigmentation of the hypocotyl are known, and this character is used in same cases used as morphological marker [43]. The QTL identified in the SABER population (LOD score peak of 29.8) is positioned at 59.0 Mb in the chromosome 9, that is inside the gene coding for a bHLH transcription factor identified as the most common responsible for absence of pigmentation in the hypocotyl [42].

Overall, we can say that all three QTL relative to mendelian traits confirm the potential of SABER as mapping population, both in accuracy and power.

### 3.4 Color of the fruit

The color of the tomato fruit at ripeness is a well-studied trait not only to address market preferences, since the bright red of the fruit or tomato-based product is one of the first traits that guide consumer choice [44], but also to unveil the complex metabolic pathways that regulate pigment accumulation in the tomato fruit. The red color of tomatoes is mainly due to carotenoids (mainly lycopene) and naringenin chalcone, which accumulate in the ripening of fruits while the chlorophyll is degraded [45]. The metabolic pathway of carotenoids has been extensively studied, as well as the genes that regulate it. *PSY1* emerged as one of the most important among them, catalyzing the first step committed in carotenoid biosynthesis. Mutants with no transcription of *PSY1* have yellow flesh [45,46]. Orange fruits are observed in a number of different mutations, resulting in intermediate lycopene synthesis and accumulation of other carotenoids (*CRTISO*, *Crtl-b*, *Crtl-e*), or limited availability of precursor of carotenoids (*IDI1*) [45,46]. The founder lines of SABER population are all with red fruits, except for the wild ancestor, *S. cheesmanie,* that has orange fruits. Among the offspring, there is a little diversification between the epicarp color (ECO: 1 yellow, 27 orange, 212 red) and the flesh color (ICO: 36 orange, 204 red). The singular yellow fruit could be a phenotypization error, since the evaluation of the color has been done with the naked eye. However, in literature it is reported a case of cross of a double mutant yellow x orange (PSY1 x CRTISO) that resulted in an orange phenotype even if the *PSY1* mutation should be dominant since the enzyme is earlier in the pathway. Kachanovsky et al. [47] found that the *CRTISO* gene could have epistatic effect on *PSY1* partially restoring its transcription. Theoretically, the yellow phenotype could emerge by segregation of the two loci from an orange individual.

The cases of different colors between flesh and epicarp were rare in the population (3 with orange epicarp and red flesh, 12 with red epicarp and orange flesh, 1 with yellow exocarp and red flesh), so it does not surprise that the QTL identified for the two phenotypes are practically identical; here they will be discussed together.

The QTL is positioned at chromosome 6, with a peak at position 43,541,536 and a confidence interval that span a physical region of 0.86 Mb (Table 3, Figure 5). The genetic determinant identified with high probability is *CYC-B* (*Solyc06g074240*, coordinates according to annotation ITAG4.1: chr6 43,562,526 – 43,564,022 bp), a lycopene-beta cyclase characterized in the tomato clade. Mohan et al. [48] demonstrated how *CYC-B* underwent purifying selection in cultivated *S. lycopersicum*, while in wild tomatoes present greater diversity in the promotorial region. Higher expression levels of *CYC-B* are associated with beta-carotene accumulation and lower levels of lycopene, resulting in orange fruits.

**Figure 5:**
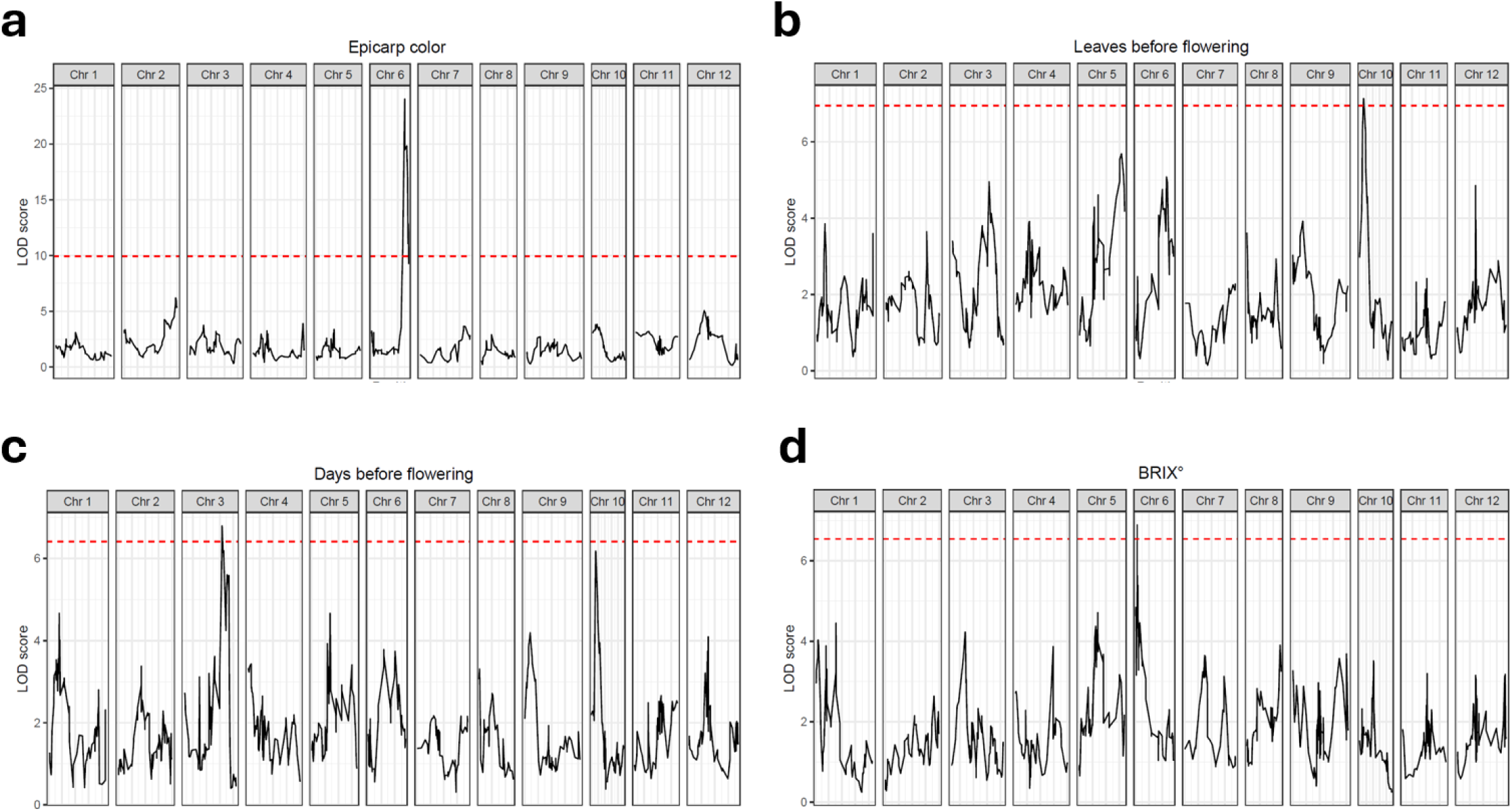
LOD scores across the genome of QTLs relative to: epicarp color (a), number of leaves before flowering (b), days before flowering (c) and BRIX value of the ripe fruit (d). Markers are positioned according to the linkage map.

**Table 3:** QTLs mapped in SABER population. Start and end position of the QTL represent the 95% Bayesian credible interval.

| Phenotype | Chromosome | Peak position (pb) | LOD score | Start (pb) | End (pb) |
| --- | --- | --- | --- | --- | --- |
| Hypocotyl color | Chr9 | 59,052,650 | 29.83 | 57,908,608 | 59,165,600 |
| Obscura venosa | Chr5 | 62,575,099 | 25.15 | 62,575,099 | 63,569,772 |
| Green shoulder | Chr10 | 2,010,429 | 8.94 | 1,208,771 | 19,083,107 |
| Epicarp color | Chr6 | 43,541,490 | 24.05 | 42,683,303 | 43,541,554 |
| Flesh color | Chr6 | 43,541,536 | 16.38 | 42,683,303 | 43,541,554 |
| BRIX° | Chr6 | 5,023,437 | 6.89 | 20,652 | 27,019,138 |
| Leaves before flowering | Chr10 | 1,738,579 | 7.13 | 1,413,363 | 1,829,496 |
| Days before flowering | Chr3 | 57,494,257 | 6.80 | 56,893,414 | 59,699,304 |

Since *CYC-B* is positioned just outside the confidence interval of the QTL, we analyzed the 122 genes inside that span (Supplementary Table X1), searching for alternative candidates. Among them, we searched for predicted genes showing homology with genes possibly involved in the external and internal coloration of the fruit. Among the most compelling candidate genes within the region, *ACO1* (1-aminocyclopropane-1-carboxylate oxidase 1; *Solyc06g073580*) clearly stands out. *ACO1* is the major *ACO* transcript accumulating during fruit ripening and catalyzes the oxidation of ACC (1-aminocyclopropane-1-carboxylic acid) to ethylene, the final step of the ethylene biosynthesis pathway [49]. Ethylene is essential for climacteric fruit ripening and plays a central role in both internal and external fruit coloration by promoting lycopene accumulation (red pigmentation) and chlorophyll degradation (green-to-red transition) [50,51]. Transcriptomic analyses suggested that ethylene production via *ACO1* influences carotenoid biosynthesis through *PSY1* expression. Conversely, reduced *ACO1* expression - such as after treatment with 1-methylcyclopropene (1-MCP) - inhibits lycopene accumulation and associated color changes, further confirming its key role in fruit pigmentation [52]. Within the same chromosomal interval, two additional and adjacent genes encode *ETHYLENE INSENSITIVE* 3 (EIN3)-like transcription factors, namely *EIL1* (*Solyc06g073720*) and *EIL4* (*Solyc06g073730*). These transcription factors are key components of the ethylene signaling cascade and may therefore also contribute to fruit pigmentation processes. Although a direct regulation of carotenoid biosynthetic genes by *EIL1* and *EIL4* has not yet been clearly demonstrated in tomatoes, evidence from other climacteric species supports their potential involvement in ripening-related pathways. In melon (*Cucumis melo*), for instance, *EIL1* has been characterized as a ripening-associated gene and shown to function as a transcriptional activator of *ACO1* [53].

### 3.5 Flowering time

Days to flowering (DTF) and leaves number before flowering (LN) are two measurements used for determining the length of the vegetative phase and are essential for the management of the plants in an industrial context. In the domestication and varieties breeding process short and uniform vegetative cycles were favored, with the goal to accelerate production and, most of all, obtain synchronous ripening of the fruits, that could be harvested at the same moment [54]. Given the importance, time flowering has been extensively documented in the literature. QTLs that influence DTF or LN have been mapped in chromosomes 1, 2, 3, 5, 6, 7, 9, and 11 [54–58] and in many cases functional characterization of the candidate genes confirmed their role [59–61]. Since the cultivated tomato shows a phenotype divergent from the wild tomatoes, in many cases the association studies have been performed on hybrid population with a wild ancestor: *S. pimpinellifolium* [56], *S. Chmielewski* [54] or *S. pennelli* [57]. This is the first study of the character *S. cheesmaniae* as parent in order to obtain variability of the phenotype.

The two measurements of the trait, the time from transplant to first flowering and the number of leaves produced are positively correlated in SABER population (Pearson correlation: 0.57, p-value < 2.2 e -16), similarly to what can be observed in literature (*ibidem*). However, the genes controlling DTF and LN are only partially overlapping, as confirmed by both QTL mapping [54] and functional studies [61]. The QTLs mapped in SABER population are consistent with the literature. The LOD peaks of the two phenotypes are positioned in two different chromosomes (Table 3), but is evident how comparing the LOD profiles there is a spot on Chr10 around 1.7 Mb with influx in both characters, even if it passes the significance threshold only for LN (Figure 4).

The QTL mapped for DTF is located on chromosome 3 from 56.8 to 59.7 Mb, a spot that that is not already known, in the literature, for its contribution to the flowering time of tomato. The LOD peak spans a 2.8 Mb region on chromosome 3 and includes 376 annotated genes (Supplementary Table X3), among which two resulted of particular interest. Foremost among these is FRUITFULL-like MADS-box 2 (*FUL2*; *Solyc03g114830*), given its well-established role in tomato in promoting the transition from vegetative to reproductive development by inducing floral meristem maturation and repressing inflorescence branching [62]. Moreover, it has been demonstrated that the *ful2 mbp20* (*Solyc02g089210*) double mutant exhibits delayed flowering [63]. Notably, the gene immediately adjacent to *FUL2*, *MADS1* (also known as ENHANCER OF JOINTLESS 2, *EJ2*; *Solyc03g114840*), has likewise been reported to influence inflorescence branching [64]. Another strong candidate within the QTL interval is SQUAMOSA PROMOTER BINDING PROTEIN 6a (*SlSBP6a*, *Solyc03g114850*), located immediately downstream of the two aforementioned genes (*FUL2* and *MADS1*), further highlighting this genomic cluster as a key regulatory hotspot for flowering-related traits. *SlSBP6a* in tomato is part of the miR156-targeted SQUAMOSA PROMOTER BINDING PROTEIN-LIKE (SPL/SBP) family, which plays a crucial role in regulating flowering time by promoting floral meristem maturation and transition to flowering. In particular, these miR156-targeted SBPs interact with gibberellin (GA) signaling pathways, synergistically regulating floral meristem determinacy and influencing flowering time [65].

For LN, the QTL confidence interval encompassed a 0.42 Mb region on chromosome 10 and included 59 annotated genes (Supplementary Table X2). Among them, we focused our attention on two consecutive genes codifying for as many Gibberellin 2-oxidases (GA2ox, *Solyc10g007570* and *Solyc10g007560*). The GA2ox family, a class of 2-oxoglutarate-dependent dioxygenases, regulates the levels of bioactive gibberellins by catalyzing their deactivation, which influences various aspects of plant growth and development [66]. While direct evidence linking GA2ox specifically to the number of leaves between cotyledons and the first truss is limited, this family broadly affects shoot architecture, leaf development, and branching patterns through modulation of gibberellin levels. For example, overexpression of GA2ox genes can lead to dwarfism, smaller leaves, and modifications in the primary and secondary growth of the stem, indicating their role in controlling vegetative growth [67–69]. Additionally, in the frame of soybean domestication, copy number variations in certain GA2ox genes were found to correlate with traits related to overall plant architecture, such as reduced shoot length and trailing growth [70].

### 3.6. Sugars and organic acids concentration in the fruits

Degrees BRIX measure the levels of dissolved solids in a solution and are the first estimate of the sugar and acids contained in tomato fruit. It is an important assessment of the quality of the product for tomatoes for every destination, industrial or fresh market. Obviously, the character has a substantial body of literature dedicated to it, and since it is a measurement of a metabolic state of the plant, it is considered a highly polygenic character, with a high number of QTLs identified. Already in 2000 Fridman et al [71] caracterized a QTL on chromosome 9, since then many more were identified in chromosomes 1, 2, 7, 8, 11 and 12 [72–76].

The QTL identified in SABER population is located in the first arm of chromosome 6 (0-27 Mb, Table 3), a position where no QTL for BRIX values was identified, with the exception of a QTL linked to the exposition of the plant to a LED light, located at 44.6 Mb [76]. The confidence interval for the position of the QTL is very broad and encompasses near 700 genes (Supplementary table X4); nevertheless, it is worth discussing some of the genes comprised in this region, considering the importance of the trait and the novelty of the identified QTL.

*Solyc06g005070* is a member of the Major Facilitator Superfamily protein (MFS), a superfamily of sugar transporters characterized by 12 transmembrane. Many members of the superfamily have been already characterized for their involvement in BRIX determination and constitute the probable genetic determinant of identified QTLs [77].

Isoamlylase 3 (*ISA3*, *Solc06g009220*) is a chloroplast enzyme deputed to the debranching of starch, identified as one of the key enzymes in starch catabolism in apple and play a role also in grape and sweet orange [78]. Starch is found only in early stages of tomato fruit [78] and its degradation could be a source for the production of sugars and organic acids.

Finally, Li et al. [79] in their analysis of BRIX-related genes using the correlation of expression level and BIX values, identified S*olyc06g005970* positively correlated with sugar levels (fructose and glucose) negatively correlated with organic acid levels (Malic and citric acid).The gene, then uncharacterized, is now annotated as a protein of the glycosyl hydrolase family

## 4. Conclusions

Expanding the genetic base of cultivated tomatoes beyond the narrow diversity retained after domestication is a central objective of contemporary breeding and genomic research. The SABER population contributes directly to this effort, being the first MAGIC population to include as a founder line an accession of *Solanum cheesmaniae,* an extremophile wild relative from the Galápagos.

The successful introgression of *S. cheesmaniae* alleles across all chromosomes, with no genomic regions showing a complete absence of wild-founder contribution, demonstrates that the SABER design effectively captured exotic genetic variation within a usable quantitative genetic framework. The relatively uniform mean founder contributions and the little population structure among the offspring further supports the population’s suitability for unbiased QTL discovery.

The testing of the potential of the population through three well-characterized Mendelian traits (obscuravenosa, uniform ripening, and hypocotyl color) confirmed the reliability of the analytical pipeline and the quality of the genotypic data obtained through SPET-based genotyping. Beyond these controls, the QTL of quantitative traits yielded both results consistent with the existing literature and novel genomic regions not previously associated with these phenotypes, suggesting that the allelic diversity introduced by the wild founder is actively contributing to phenotypic variation within the population.

From a broader perspective, SABER establishes a replicable model for integrating wild germplasm into multiparental breeding resources. As climate change increasingly threatens the productivity and resilience of major crops, the ability to exploit the adaptive potential encoded in wild relatives becomes strategically essential. Future work should focus on expanding phenotyping efforts — particularly for abiotic stress-related traits — and on fine-mapping the novel QTL regions identified here to accelerate candidate gene characterization and marker-assisted selection.

## Data availability

The datasets generated and analyzed during the current study are available in National Center for Biotechnology Information (NCBI) and can be accessed in the sequence read archive (SRA) database (https://www.ncbi.nlm.nih.gov/sra). The accession number is PRJNA1417763.

## List of supplementary data

### Supplementary figures

**Figure S1** – Density of SNPs with genotypes alternative to the reference SL4.0 of the eight founder lines, plotted binning the genomes in 200Kb windows. The twelve vertical bars represent the twelve chromosomes, the eight founder lines are represented by the colored box under each cromosomes. The coordinates of the chromosomes are in the y-axis.

### Supplementary tables

**Table S1** – Phenotypes of F6 samples. Hypocotyl color (ANT = purple, AH = green), Obscuravenosa (NT = not transparent, T = transparent, Green shoulder (G = present, UG = not present), Epicarp and Flesh color (R = red, Beta = orange, Y = yellow). The quantitative traits (°Brix values, leaves before flowerin LN, Days to flowering DTF) are measured as descripted in chapter 2.4.

**Table S2** – Pedigree and proprieties of the eight founder lines

**Table S3** – Linkage map of SABER with the physical (bp) and genetic (cM) position of each marker

**Table S4** – Sequencing, alignment and SNP calling metrics. The columns contain: sample ID, number of pairs of total unfiltered reads (total sequenced reads), number of pairs of filtered reads (filtered reads), number of successfully aligned reads (aligned reads), number of reads aligned with a quality equal or higher than 5 (aligned reads q5), average depth of reads on SNP called (Average depth), number and percentage of: loci with homozygous genotype equal to the reference genome SL4.0 (Reference homozygous), loci with homozygous genotype alternative to the reference genome SL4.0 (Alternative Homozygous), heterozygous loci (Heterozygous) and loci with missing genotypes (Missing)

**Table S5** – List of genes and their functional annotation comprised in LOD curve of QTL for epicarp and flesh color, according to annotation ITAG4.1

**Table S6** - List of genes and their functional annotation comprised in LOD curve of QTL for DTF, according to annotation ITAG4.1

**Table S7** - List of genes and their functional annotation comprised in LOD curve of QTL LN, according to annotation ITAG4.1

**Table S8** - List of genes and their functional annotation comprised in LOD curve of QTL for °Brix, according to annotation ITAG4.1

### Supplementary files

**Data S1** – chain files to lift over the coordinates from *S. lycopersicum* annotation SL2.4 to SL4.0

## Notes

### Competing Interest Statement

The authors have declared no competing interest.

