## Supplementary figures and images for "SABER: A Multiparental Tomato Population Leveraging Wild Relative Diversity for High-Resolution QTL Mapping"

### Figure S1

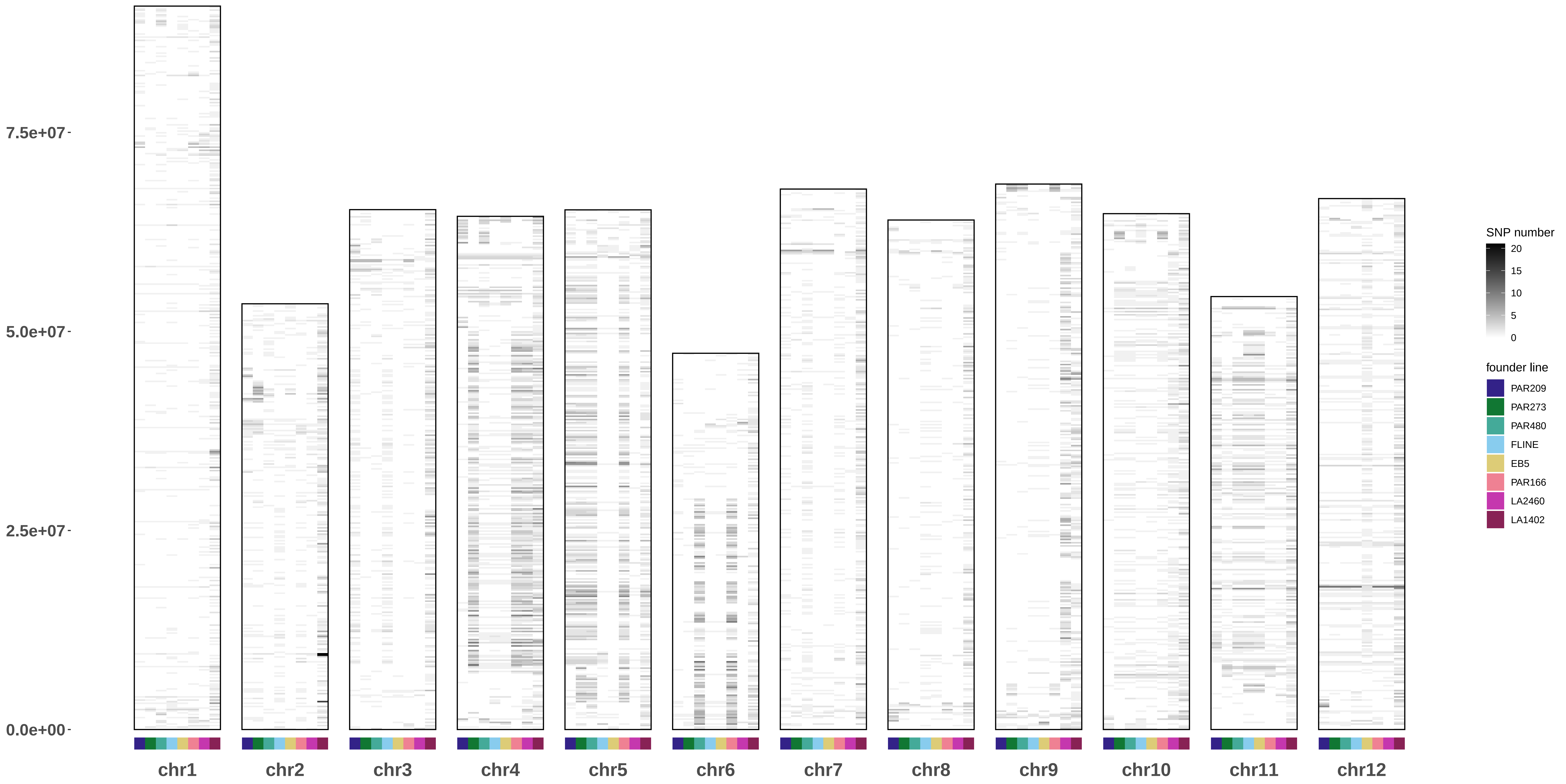
